# Impact of Serum Circulating Factors and PDE5 Inhibitor Therapy on Cardiomyocyte Metabolism in Single Ventricle Heart Disease

**DOI:** 10.1101/2025.03.31.646497

**Authors:** Anastacia M. Garcia, Ashley E. Pietra, Mary E. Turner, Julie Pires Da Silva, Angela N. Baybayon-Grandgeorge, Genevieve C. Sparagna, Danielle A. Jeffrey, Brian L. Stauffer, Carmen C. Sucharov, Shelley D. Miyamoto

## Abstract

**Background:** While operative and perioperative care continues to improve for single ventricle congenital heart disease (SV), long-term morbidities and mortality remain high. Importantly, phosphodiesterase-5 inhibitor therapies (PDE5i) are increasingly used, however, little is known regarding the direct myocardial effects of PDE5i therapy in the SV population.

**Objectives:** Our group has previously demonstrated that the failing SV myocardium is characterized by increased PDE5 activity and impaired mitochondrial bioenergetics. Here we sought to determine whether serum circulating factors contribute to pathological metabolic remodeling in SV, and whether PDE5i therapy abrogates these changes.

**Methods:** Using an established *in vitro* model whereby primary cardiomyocytes are treated with patient sera +/− PDE5i, we assessed the impact of circulating factors on cardiomyocyte metabolism. Mass spectrometry-based lipidomics and metabolomics were performed to identify phospholipid and metabolite changes. Mitochondrial bioenergetics were assessed using the Seahorse Bioanalyzer and a stable isotope based mitochondrial enzyme activity assay. Relative mitochondrial copy number was quantified using RT-qPCR.

**Results:** Our data suggest that serum circulating factors contribute to fundamental changes in cardiomyocyte bioenergetics, including impaired mitochondrial function associated with decreased cardiolipin and other phospholipid species, increased reactive oxygen species (ROS) generation, and altered metabolite milieu. Treatment with PDE5i therapy was sufficient to abrogate a number of these metabolic changes, including a rescue of phosphatidylglycerol levels, a reduction in ROS, improved energy production, and normalization of several key metabolic intermediates.

**Conclusions:** Together, these data suggest PDE5i therapy has direct cardiomyocyte effects and contributes to beneficial cardiomyocyte metabolic remodeling in SV failure.

## 1. Introduction

Single Ventricle congenital heart disease (SV) encompasses a myriad of congenital cardiac malformations and is typified by the severe underdevelopment of one ventricle[1]. Morphologically, SV can be characterized by the underdevelopment of either the right (RV) or left ventricle (LV), however, patients with a systemic RV, such as those with hypoplastic left heart syndrome (HLHS), represent the most common SV subtype and tend to have worse outcomes[2–7]. However, while outcomes are improving, SV remains universally fatal without intervention, and greater than 30% of SV patients die or require transplant within the first year of life [8, 9]. Heart failure (HF) is the leading cause of death and indication for transplant in SV [9], and despite advancements in perioperative and surgical care, degradation of RV function and eventual heart failure still significantly impact long-term survival and quality of life for these patients. However, the molecular mechanisms underlying single RV failure remain poorly understood, partially due to the lack of a postnatal animal model of SV and limitations of *in vivo* pediatric research. As such, the current standards of care for SV HF are based on expert consensus and the extrapolation of adult HF therapies, though outcomes for children with SV have not improved to nearly the same extent as adults when given standard HF therapies [10, 11]. In particular, various pediatric clinical trials of commonly utilized adult HF therapeutics, such as carvedilol and enalapril, have demonstrated a lack of response [12] or a detrimental response [13] in preventing or rescuing SV HF.

Importantly, while there are no proven medical therapies for the treatment or prevention of HF in the SV population, selective and competitive phosphodiesterase-5 inhibitors (PDE5i), such as sildenafil, tadalafil, and udenafil, are increasingly utilized. Nevertheless, while there is mounting evidence suggesting beneficial effects of PDE5i therapy on exercise tolerance and hemodynamics in the SV population, presumably related to the pulmonary vasodilator effects of these medications, little is known regarding any direct myocardial effects [14–17]. The most recent multicenter, double-blind, randomized, placebo controlled FUEL (Fontan Udenafil Exercise Longitudinal) trial, which assessed the exercise capacity and ventricular function of stage-3 palliated (post-Fontan) SV patients treated with long acting PDE5i, indicated that while treatment with udenafil (87.5 mg twice daily) was not associated with an improvement in oxygen consumption at peak exercise, it was associated with improvements in multiple measures of exercise performance at the ventilatory anaerobic threshold, particularly in patients with a systemic RV [18–20]. Additionally, while udenafil did not significantly change systolic function, FUEL participants who received udenafil demonstrated a statistically significant improvement in some global and diastolic echo indices (e.g., myocardial performance index, atrioventricular valve inflow peak E and A velocities, and annular Doppler tissue imaging-derived peak e’ velocity) [18–20]. It is important to note that the Fontan patients included in this trial had relatively normal single ventricle function (normal ventricular ejection fractions) and exercise tolerance, therefore, it remains unclear whether PDE5i therapy could be more clinically beneficial in SV patients with significantly impaired exercise tolerance and or medically refractory HF. Nevertheless, these data suggest that PDE5i therapy had a small but measurable impact on SV mechanics, which may reflect a direct influence on the myocardium.

We have previously demonstrated that PDE5 expression and activity are significantly elevated in failing SV myocardium, further suggesting potential for direct myocardial effects of PDE5i therapy. Additionally, using an established novel *in vitro* model consisting of treating primary cardiomyocytes with serum from pediatric SV HF patients as compared to those treated with serum from healthy age-matched controls, we determined that PDE5i is sufficient to abrogate SV serum-induced pathological gene expression changes *in vitro* [21]. In the present study, using the same *in vitro* model, we sought to further define potential direct cardiomyocyte effects of PDE5i therapy. We determined that SV circulating factors promote metabolic remodeling and mitochondrial dysfunction in primary cardiomyocytes, and that many of these pathological bioenergetic changes are abrogated with sildenafil treatment, suggesting the possibility of direct beneficial myocardial metabolic effects of PDE5i.

## 2. Methods

### 2.1 Human Samples

The studies performed were conducted according to Declaration of Helsinki principles and subjects or guardians of subjects under the age of 18 years gave written informed consent prior to inclusion in the study. The study was approved by the University of Colorado Anschutz Medical Campus Institutional Review Board. Samples of non-failing control sera were obtained from healthy children (<18 years of age) with normal cardiac structure and function (Healthy Ctrl). Failing SV (SV HF) sera were obtained from patients with SV of RV morphology, <18 years of age, who had decreased right ventricular systolic function on transthoracic echocardiogram, medically refractory protein-losing enteropathy (PLE), plastic bronchitis (PB), and/or right ventricular end-diastolic pressure or pulmonary capillary wedge pressure of ≥12 mm Hg. Determination of systolic function was made on the basis of review of the clinical echocardiographic report for each subject, patients that were transplanted primarily for SV lacking these defined clinical characteristics were excluded. Patients with single ventricle disease of LV or indeterminate morphology were also excluded.

### 2.2 Serum Separation

Whole blood was drawn from patients and stored at 4°C in a red top BD Vacutainer Blood Collection Tube with clot activator silicone coating. The red top tube sat at room temperature for 30 minutes before centrifugation. It was spun at 4°C at 600 x g for 20 minutes in a swinging bucket rotor centrifuge. The serum layer was transferred from the red top tube to 1.5mL Eppendorf tubes and stored at –80°C until further use.

### 2.3 Primary Cardiomyocyte Cell Culture and Treatments

Neonatal Rat Ventricular Myocytes (NRVMs) were isolated from 1-2-day-old Sprague-Dawley rats (Charles River) by enzymatic digestions as described [21]. NRVMs were incubated with 1.5 µmol/L sildenafil for 30 minutes before serum treatment. NRVMs were then treated with a final concentration of 2% serum and left to incubate for 72 hours before further experiments. All animal protocols are in accordance with Public Health Service Animal Welfare Assurance, ID A3269-01, and approved by the University of Colorado, Denver – Animal Care and Use Committee.

### 2.4 Phospholipid Quantification

Phospholipids were extracted from NRVM samples that were washed and harvested in PBS. Phospholipids were quantified using LC coupled to electrospray ionization MS in an API 4000 mass spectrometer, as described [22]. Tetramyristoyl cardiolipin (1,000 nmol) and Splash phospholipid standards (Avanti Polar Lipids) were used as internal reference standards to identify phospholipid retention times. Phospholipid species were quantified per milligram of protein on the basis of a protein assay of the NRVMs (BioRad Bradford Quickstart Assay Kit). Analysis was performed using Analyst software and exported data were expressed in fold change relative to controls within each prep. Summed phospholipids included the following species: cardiolipin (CL) sum of mass/charge (m/z) 1422, 1448, 1450, 1452, 1470, 1472, 1496 and 1498; phosphatidylcholine (PC) sum of m/z 818, 840, 844, and 868; phosphatidylethanolamine (PE) sum of m/z 714, 716, 738, 742, 762, 764, and 766; phosphatidylglycerol (PG) sum of m/z 747, 771, and 773; phosphatidylinositol (PI) sum of m/z 857, 861, 883, 885, 909, and 911. The m/z ratio and fatty acyl chain identification for each of the major CL species shown is listed in **Table S2**.

### 2.5 Mitochondrial Bioenergetics Analysis

NRVMs were plated on a sterile gelatin-coated 96-well Seahorse plate at a density of 60,000 cells per well. Cells were treated in replicates of 3-5 wells per condition, with sildenafil and serum, as discussed above. The Cell Mito Stress Test was used with an Agilent XFe 96-well Seahorse Bioanalyzer to quantify standard oxidative phosphorylation (OXPHOS) or FAO. The day prior to the assay, a calibration cartridge was hydrated with sterile Seahorse calibrant solution at 37°C in a non-CO_2_ incubator overnight. For the fatty acid oxidation assay (FAO), cells were treated with 5mM L-carnitine for 24-hours prior to the assay. One hour before the standard OXPHOS assay, the cells were gently washed and supplied with warmed seahorse media (DMEM plus substrates for a final concentration of 1mM pyruvate, 2mM glutamine, and 10mM glucose, pH to 7.4 with NaOH), and cells were incubated for at least 45 min at 37°C in a non-CO_2_ incubator to de-gas.

One hour before the FAO assay the cells were gently washed and supplied with warmed FAO media (DMEM plus 0.5mM L-carnitine and 5mM HEPES), and cells were incubated for at least 45 min at 37°C in a non-CO_2_ incubator to de-gas. To quantify FAO specifically, during the 45min incubation, a subset of cells were treated with either 45uM etomoxir (negative control) or vehicle, and with BSA-conjugated LCFA palmitate (FAO). Freshly diluted inhibitors were pre-loaded to the calibration cartridge in three separate ports for a final well concentration of: 1.5µM oligomycin, 1.0 µM carbonyl cyanide-4-(trifluoromethoxy) phenylhydrazone (FCCP), and 0.9 µM each of rotenone/antimycin A. After the Seahorse assay, data were normalized between wells using a CyQUANT™ Direct Cell Proliferation Assay (ThermoFisher Scientific) as described [23] to quantify the number of live cells per well. Cells were incubated for an hour in CyQUANT dye and imaged with a fluorescent plate reader with excitation set to 485/20 and emission 528/20. Mitochondrial oxygen consumption rate (OCR) was calculated as pmol/min/live cells after normalization. All parameters of mitochondrial bioenergetics were calculated according to equations listed in **Table 2**.

### 2.6 Reactive Oxygen Species (ROS) Quantification

Superoxide and hydrogen peroxide were measured together in cells using Amplex UltraRed Reagent assay kit (ThermoFisher Scientific) as described [24]. Cells were plated in a black, clear-bottom 96-well microplate and treated with serum and Sildenafil as described previously, 72 hours prior to running the assay. After 72 hours, the cells were washed in 100µL of PBS. A solution made up of Amplex red dye (final concentration: 50µM), hydrogen peroxide (0.0015% final), +/− superoxide dismutase (SOD, final concentration: 5 units/mL) was added to each well to bring the final volume to 200µL. The plate was then incubated, in the dark, at 37°C for 30 minutes. Sample fluorescence was measured using a SynergyHTX Mutli-Mode Plate Reader (BioTek) at maximum excitation and emission wavelengths of 540 and 600 nm, respectively. The plate was then washed twice with 100µL of PBS and live cell number was assessed using CyQUANT cell proliferation assay kit according to manufacturer’s instructions (ThermoFisher Scientific). Each condition was examined at least in quadruplicate (4-5 wells/condition). Data were calculated as average values of ROS per live cell count and expressed in fold change relative to controls within each prep.

### 2.7 Relative Mitochondrial Copy Number Quantification

DNA was extracted using the DNeasy Blood and Tissue kit (Qiagen) and quantified with Quant-iT^TM^ PicoGreen dsDNA kits (Invitrogen) as per manufacturer’s instructions. RT-qPCR of nuclear and mitochondrial DNA sequences was performed to assess the proportion of mitochondrial genomes relative to nuclear DNA as a surrogate for mitochondrial content. Mitochondrial DNA (mtDNA) sequences MT-ND5 and MT-CytB were normalized to nuclear β2-microglobulin (B2M) [25].

### 2.8 Metabolomics Analysis

Untargeted metabolomics analysis was performed by University of Colorado Metabolomics Core Facility by using ultrahigh-pressure (UP) liquid chromatography (LC; UPLC) coupled to online mass spectrometry (MS). Briefly, NRVM sample extracts were injected into a UPLC system (Ultimate 3000, Thermo) and run on a Kinetex XB-C18 column (150 × 2.1 mm and 1.7-μm particle size, Phenomenex) at 250 μL/min (mobile phase, 5% acetonitrile, Sigma-Aldrich), 95% 18 mΩ H2O (Sigma-Aldrich), and 0.1% formic acid (Sigma-Aldrich). The UPLC system was coupled online with a Q Exactive system (Thermo), scanning in Full MS mode (2 microscans) at a 70,000 resolution in the 60-to 900-m/z range, 4-kV spray voltage, 15 sheath gases and 5 auxiliary gases, operated in negative-ion and then positive-ion mode (separate runs). Calibration was performed before each analysis against positive-ion or negative-ion mode calibration mixes (Pierce, Thermo Fisher) to ensure subparts per million error of the intact mass. Metabolite assignments were performed using Maven software,32 on conversion of .raw files into .mzXML format through MassMatrix software. Such software allows peak picking, feature detection, and metabolite assignment against the KEGG (Kyoto Encyclopedia of Genes and Genomes) pathway database to be conducted. Assignments were further confirmed against chemical formula determination (as gleaned from isotopic patterns and accurate intact mass), and retention times were confirmed against an in-house validated standard library (>650 compounds, including various metabolites, amino acids [AAs], and acylcarnitines) (Sigma-Aldrich; MLSMS, IROATech). Relative quantitation was performed by exporting integrated peak area values into Excel (Microsoft) for statistical analyses. Hierarchical clustering, enrichment, and pathway analysis was conducted using MetaboAnalyst 6.0.

### 2.9 Carnitine Palmitoyltransferase Enzymatic Activity

Carnitine palmitoyltransferase (CPT) I and CPTII enzyme activities were quantified in serum-treated NRVMs using a carbon-14 (^14^C) labeled carnitine radioactivity assay as described [22, 26]. Briefly, CPTI activity was measured by permeabilizing the plasma membrane and quantifying the production of ^14^C labeled palmitoylcarnitine from palmitoyl-CoA. The activity of CPT II was measured by permeabilizing the inner mitochondrial membrane and adding malonyl CoA to inhibit CPT I.

### 2.10 Statistical Analysis

Statistical analyses were performed with GraphPad Prism version 10.2.2, and statistical significance was set a priori at P < 0.05. All n values, statistical tests used, and descriptions of graphical depictions of data are defined within the figure legends for the respective data panels. Data were tested for gaussian distribution with the Shapiro-Wilk normality test. Welch’s correction was used when variances were unequal on the basis of the F-test. Comparisons between 2 normally distributed groups were conducted using an unpaired t-test, comparisons between 2 non-normally distributed groups were conducted using the nonparametric Mann-Whitney test, and comparisons between 2 groups with unequal variances were conducted using Welch’s t-test. Comparisons of 3 or more normally distributed groups were conducted using an oridinary one-way ANOVA; if the overall comparison reached significance, a Holm-Sidak post hoc test for multiple pairwise comparisons was performed. Comparisons among 3 or more non-normally distributed groups were conducted using the nonparametric Kruskal-Wallis test; if the overall comparison reached significance, Dunn’s post hoc test for multiple pairwise comparisons was performed. Comparisons among 3 or more groups with unequal variances were conducted using Welch’s ANOVA; if the overall comparison reached significance, Dunnett’s post hoc test for multiple pairwise comparisons was performed.

## 3. Results

### 3.1 Patient Characteristics

Aggregate characteristics for patients included in this study are listed in **Table 1**. Included in Supplemental Table 1 (**Table S1**) is a more detailed description of the individual patient characteristics. The healthy control group included patient serum from 29 subjects with a median age of 8.7 years, an interquartile range of 6.1 to 14, and was 62.07% male. The SV HF group included patient serum from 35 subjects with a median age of 3.9 years, an interquartile range of 2.97-9.7, and was 68.57% male. Due to the limited availability of young healthy controls, the SV HF patient group is slightly younger than the healthy control group (P= 0.0502).

**Table 1.** Patient Characteristics Aggregate characteristics for all patients included in this study. SVHF = single-ventricle heart failure, IQR=interquartile range, ROS = reactive oxygen species, mtDNA = mitochondrial DNA copy number, CPT = carnitine palmitoyltransferase.

| Experiment | Group | % Male | Median Age (years) [IQR] |
| --- | --- | --- | --- |
| Lipidomics | Healthy | 75% | 5.5 [5.075-7.35] |
|  | SVHF | 71% | 2.8 [2.0-9.3] |
| ROS | Healthy | 69% | 12.6 [5.4 - 16.3] |
|  | SVHF | 50% | 4.75 [4.1-8.775] |
| Bioenergetics | Healthy | 80% | 5.4 [4.1-12.6] |
|  | SVHF | 86% | 4.2 [1.65-8.4] |
| Metabolomics | Healthy | 67% | 5.4 [4.75-11.45] |
|  | SVHF | 67% | 4.8 [4.5-7.45] |
| mtDNA | Healthy | 75% | 5.5 [5.075-7.35] |
|  | SVHF | 80% | 6.7 [4.2-14.8] |
| CPT Activity | Healthy | 50% | 8.15 [6.425-13.775] |
|  | SVHF | 86% | 8.1 [4.4-14.4] |

**Table 2.** Agilent Seahorse XF Cell Mito Stress. Test Equations Equations used to calculate various parameters of mitochondrial bioenergetics using the combination of Oligomycin, FCCP, and Rotenone/Antimycin A to conduct the Cell Mito Stress Test using the Agilent Seahorse Bioanalyzer

| Parameter Value | Equation |
| --- | --- |
| **Non-Mitochondrial Oxygen Consumption** | Minimum rate measurement after Rotenone/Antimycin A injection |
| **Basal Respiration** | (Last rate measurement before first injection) – (Non-Mitochondrial Respiration) |
| **Maximal Respiration** | (Maximum rate measurement after FCCP injection) – (Non-Mitochondrial Respiration) |
| **Proton (H<sup>+</sup>) Leak** | (Minimum measurement after Oligomycin injection) – (Non-Mitochondrial Respiration) |
| **ATP Production** | (Last rate measurement before Oligomycin injection) – (Minimum rate measurement after Oligomycin injection) |
| **Spare Respiratory Capacity** | (Maximal Respiration) – (Basal Respiration) |
| **Coupling Efficiency (%)** | (ATP Production) / (Basal Respiration) X 100 |
| **Bioenergetic Health Index** | $\log \frac{(\text{Spare Respiratory Capacity}) \times (\text{ATP Production})}{(\text{Non-Mitochondrial Oxygen Consumption}) \times (\text{Proton Leak})}$ |

### 3.2 Failing SV Serum Decreases Cardiomyocyte Phospholipid Content

To determine whether cardiomyocyte phospholipid abundance is altered in response to SV circulating factors, mass spectrometry based lipidomic analysis was performed on serum treated primary cardiomyocytes (**Figure 1**). Total content of each phospholipid molecular species was quantified in nanomoles per milligram of protein. Total mitochondrial cardiolipin (CL) content is significantly decreased (P=0.0147) in NRVMs treated with SV HF patient serum relative to those treated with healthy control serum. Additionally, the quantity of several other phospholipid species was significantly decreased in response to SVHF serum treatment, including phosphatidylcholine (PC, P=0.0097); phosphatidylethanolamine (PE, P=0.0007), phosphatidylglycerol (PG, P=0.0006), and phosphatidylinositol (PI, P=0.0010). Given the major role CL plays in mitochondrial structure and function, each of the individual major CL species were also quantified. The most abundant form of CL in the heart, tetralinoleoyl CL (m/z 1448), is significantly decreased (P=0.0157) in SVHF serum-treated cardiomyocytes (P=0.0157), as are CL 1422 (P=0.0177), CL 1472 (P=0.001), CL 1496 (P=0.0190), and CL 1498 (P=0.0006) species.

**Figure 1.**
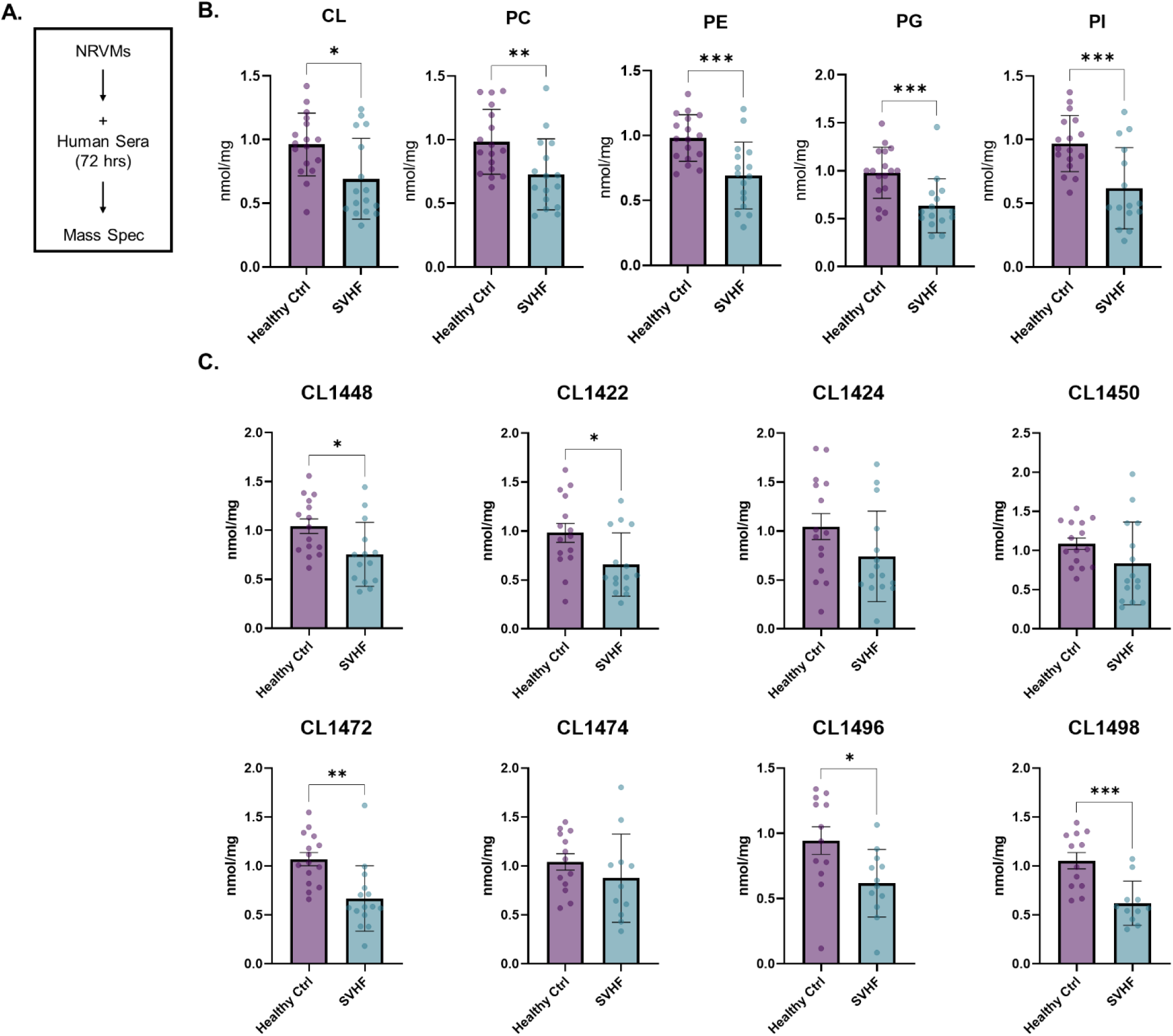
Failing SV Serum Decreases Cardiomyocyte Phospholipids *in vitro*. A: Mass spectrometry–based phospholipid analysis of phospholipid species in serum-treated primary cardiomyocytes. B: Quantification (normalized nmol/mg) of each detected phospholipids species including cardiolipin (CL), phosphatidylcholine (PC), phosphatidylethanolamine (PE), phosphatidylglycerol (PG), phosphatidylinositol (PI). C: Quantification (normalized nmol/mg) of specific CL species. Bar equals mean ± SEM; each dot represents an individual patient serum-treated NRVM: n=4 healthy controls and n=7 single ventricle heart failure samples; asterisks denote significant differences among groups; ∗P <0.05, ∗∗P<0.01, and ∗∗∗P <0.0001; analysis using Mann-Whitney test for CL and PG and Unpaired T-test for PC, PE, and PI, and all individual CL species.

### 3.3. Failing SV Serum Increases Reactive Oxygen Species Generation and Alters Cardiomyocyte Mitochondrial Bioenergetics

To assess whether serum-induced alterations in phospholipid content impair mitochondrial function, we conducted the Cell Mito Stress Test in the Agilent Seahorse Bioanalyzer (**Figure 2**). Treatment of primary cardiomyocytes with failing SV patient sera resulted in significantly decreased basal respiration (P=0.0073), ATP Production (P=0.0138), coupling efficiency (P=0.0351), and the overall Bioenergetic Health Index (P=0.0170). There were no differences in maximal respiration, spare respiratory capacity, proton leak, or non-mitochondrial oxygen consumption. Additionally, we quantified the production of reactive oxygen species (ROS), and determined that SV HF patient sera is sufficient to significantly increase cardiomyocyte ROS production (P=0.0112).

**Figure 2.**
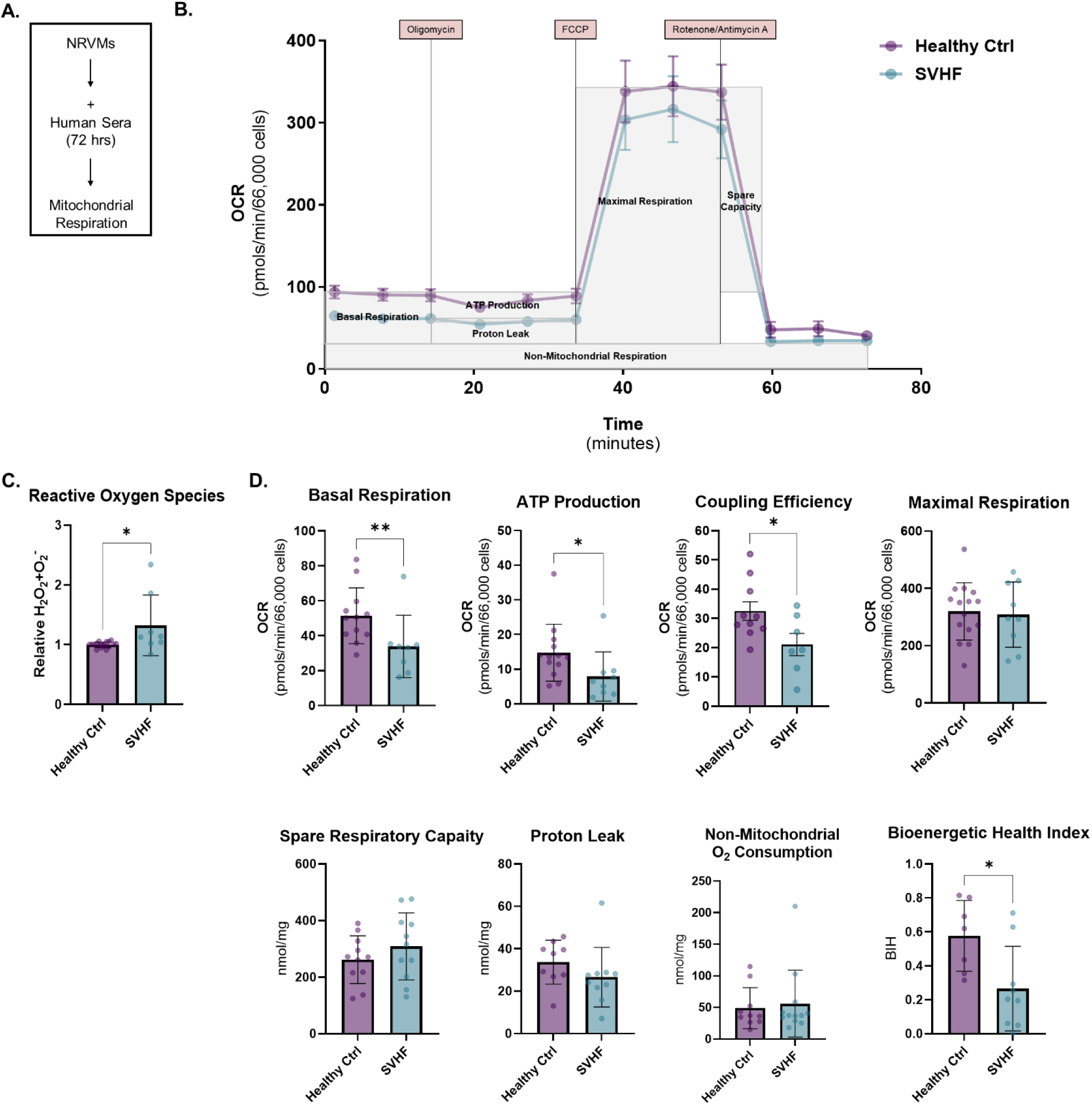
Failing SV Serum Increases Reactive Oxygen Species Generation and Alters Cardiomyocyte Mitochondrial Bioenergetics *in vitro*. A: Mitochondrial respiration in serum –treated primary cardiomyocytes. B: Representative trace of oxygen consumption rate (OCR) curve in NRVMs. Cells were exposed sequentially to oligomycin, carbonyl cyanide p-(tri-fluromethoxy)phenyl-hydrazone (FCCP) and rotenone/antimycin (AA). C: Quantification (relative H_2_O_2_+O ^-^) of Reactive Oxygen Species (ROS); Bar equals mean ± SEM; each dot represents an individual patient serum-treated NRVM: n=13 (6 patient replicates) Healthy Controls and n=6 (2 patient replicates) failing SV serum treated NRVMs across a total of 10 NRVM preparations; asterisk denotes significant difference among groups; ∗P <0.05; analysis using Unpaired T-test. D: Respiratory parameters such as basal respiration, ATP production, coupling efficiency, maximal respiration, spare respiratory capacity, proton leak, non-mitochondrial oxygen consumption, and bioenergetic health index were calculated. Bar equals mean ± SEM; each dot represents an individual serum treated NRVM sample: n=12 healthy controls, n=8 SVHF samples; asterisks denote significant differences among groups; ∗P <0.05 and ∗∗P<0.01; analysis using Unpaired T-test for coupling efficiency and maximal respiration, Mann-Whitney test for basal respiration, ATP production, spare respiratory capacity, proton leak, and non-mitochondrial oxygen consumption, and Unpaired T-test with Welch’s correction for bioenergetic health index.

### 3.4 Failing SV Serum Induces Metabolite Changes in Cardiomyocytes

To assess whether circulating factors are sufficient to alter the metabolic signature in primary cardiomyocytes, global metabolomics analysis was conducted. Metabolomics analysis identified more than 100 metabolites in NRVMs, 19 of which were significantly differentially expressed (P<0.05) in response to SVHF serum treatment as compared to healthy control serum treatment (**Figure 3**). Unsupervised hierarchical clustering of the top 30 differentially regulated metabolites separated healthy control and SVHF serum-treated cells. Pathway analysis of the 19 significantly differentially expressed metabolites using Metaboanalyst revealed enrichment of pathways involved in purine metabolism, linoleic acid metabolism, long-chain fatty acid β-oxidation, and glycolysis.

**Figure 3.**
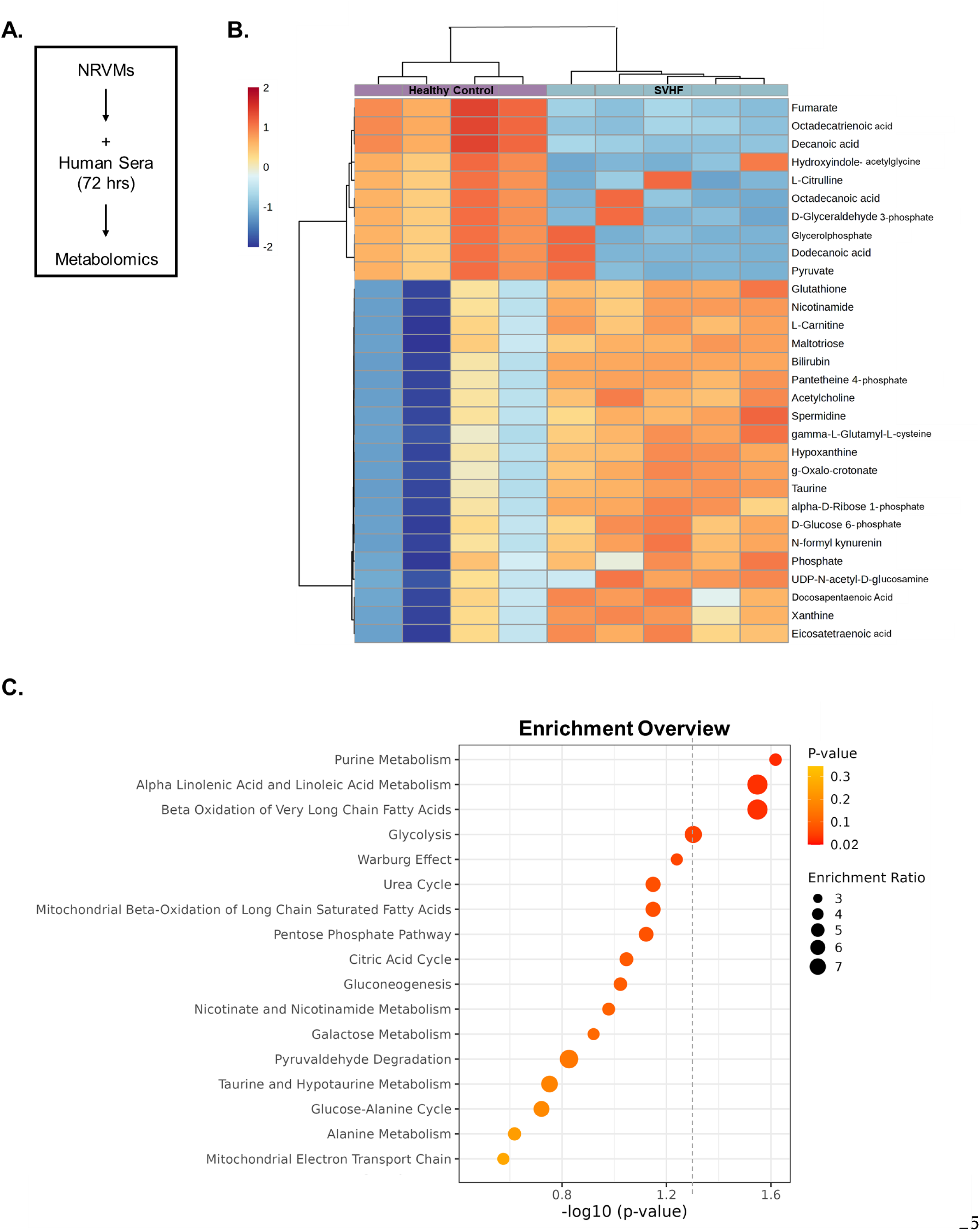
Failing SV Serum Induces Metabolite Changes in Cardiomyocytes in vitro. A: Mass spectrometry–based metabolomics analysis in serum-treated primary cardiomyocytes. B: Heatmap representation of the top 30 differentially expressed metabolites in serum-treated NRVMs. Unsupervised hierarchical clustering of the top 30 metabolites separated healthy control and failing SV serum-treated cells. C: Significantly enriched canonical pathways identified with Metaboanalyst using the 19 metabolites that changed significantly in the SV HF sera-treated NRVMs; Fisher’s exact test, – log_10_ (P-value) >1.3 or P<0.05. For all groups, average of n=4 (1 patient replicate) Healthy Controls and n=5 (2 patient replicates) failing SV patient-serum treated NRVMs across a total of 5 NRVM preparations.

### 3.5 PDE5i Does Not Attenuate Most SV Serum-Induced Phospholipid Changes in Cardiomyocytes

Using the same serum-based *in vitro* model of cardiomyocyte remodeling, we assessed the intracellular consequences of PDE5 inhibition on overall metabolic function. While SVHF sera was sufficient to significantly decrease total cardiomyocyte phospholipid levels, PDE5i treatment did not significantly abrogate the changes in CL, PC, PE, or PI in response to SV HF sera (**Figure 4**). However, PDE5i treatment did significantly increase PG content in SVHF serum-treated cells (P=0.0259).

**Figure 4.**
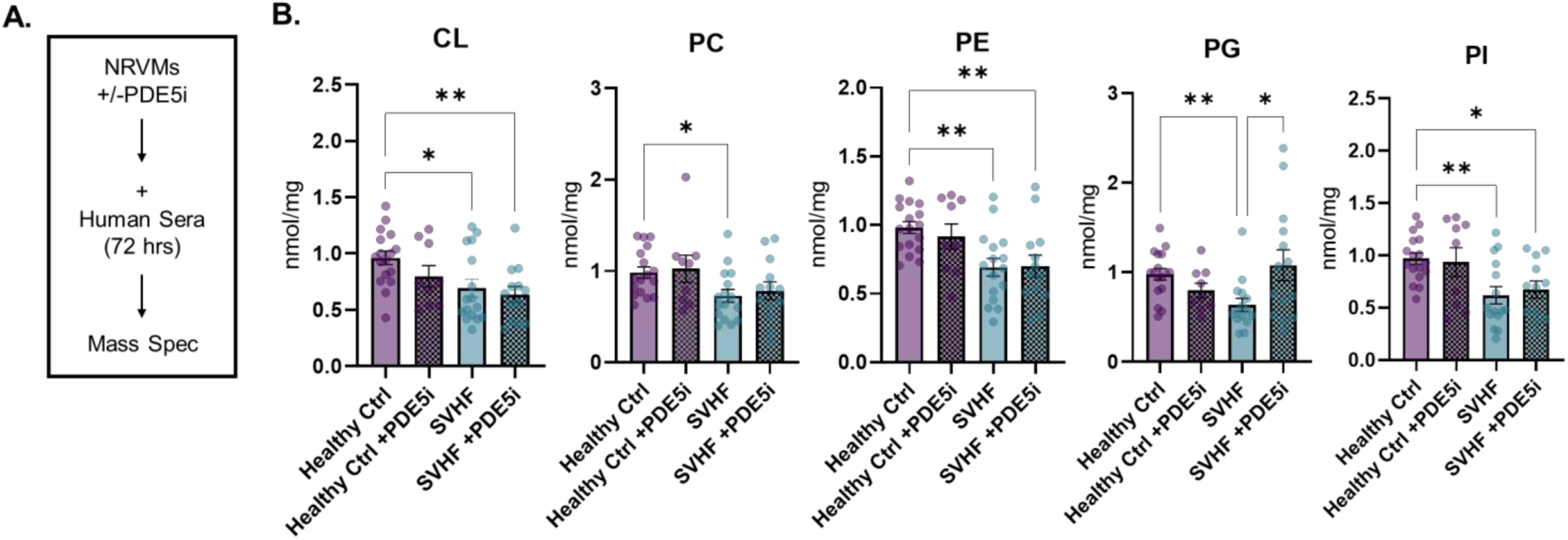
PDE5i Does Not Attenuate Most SV Serum-Induced Phospholipid Changes in Cardiomyocytes *in vitro*. A: Mass spectrometry–based phospholipid analysis of various phospholipid species in serum-treated primary cardiomyocytes +/− PDE5i treatment. B: Quantification (nmol/mg) of each detected phospholipids species; cardiolipin (CL), phosphatidylcholine (PC), phosphatidylethanolamine (PE), phosphatidylglycerol (PG), phosphatidylinositol (PI) with and without PDE5i. Bar equals mean ± SEM; each dot represents an individual patient serum-treated NRVM: n=13 (6 patient replicates) Healthy Controls and n=6 (2 patient replicates) failing SV serum treated NRVMs +/− PDE5i across a total of 10 NRVM preparations; asterisks denote significant differences among groups; ∗P <0.05 and ∗∗P<0.01; analysis using Kruskal-Wallis and post hoc Dunn’s multiple comparisons test for CL, PG, and PI and Ordinary one-way ANOVA and post hoc Holm-Sidak’s multiple comparisons for PC and PE.

### 3.6 PDE5i Attenuates SV Serum-Induced Mitochondrial Bioenergetic Changes in Cardiomyocytes

To further investigate whether PDE5 inhibitor treatment modifies mitochondrial function, we assessed oxygen consumption rates (OCR) of serum-treated primary cardiomyocytes +/− PDE5i using the same Cell Mito Stress Test in the Agilent Seahorse Bioanalyzer. PDE5i treatment was sufficient to rescue pathological changes in various parameters of mitochondrial function (**Figure 5**). Specifically, we observed that PDE5i treatment significantly abrogated the SVHF serum induced pathological changes in ATP production and coupling efficiency, and was sufficient to significantly decrease proton leak (P=0.025) and significantly increase the overall bioenergetic health index (P=0.0253). Moreover, PDE5i therapy significantly reduced ROS production in SVHF serum-treated cardiomyocytes (P=0.0382).

**Figure 5.**
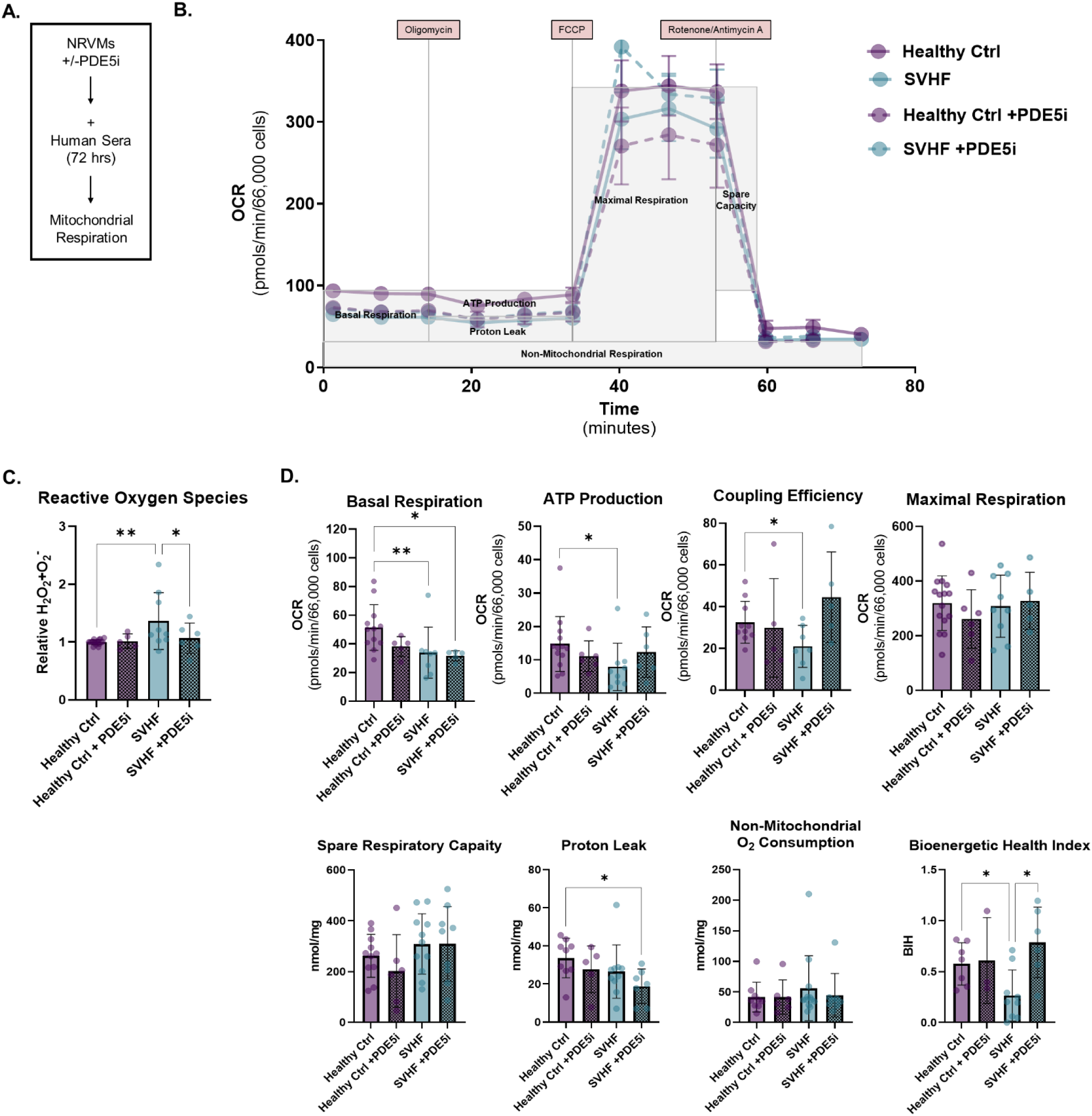
PDE5i Attenuates SV Serum-Induced Mitochondrial Bioenergetic Changes in Cardiomyocytes in vitro. A: Mitochondrial respiration in serum-treated primary cardiomyocytes +/− PDE5i. B: Representative trace of oxygen consumption rate (OCR) curve in NRVMs. Cells were exposed sequentially to oligomycin, carbonyl cyanide p-(tri-fluromethoxy)phenyl-hydrazone (FCCP) and rotenone/antimycin (AA). C: Quantification (relative H_2_O_2_+O ^-^) of Reactive Oxygen Species (ROS). Bar equals mean ± SEM; each dot represents an individual patient serum-treated NRVM: n=13 (6 patient replicates) Healthy Controls and n=6 (3 patient replicates) failing SV patient-serum treated NRVMs +/− PDE5i across a total of 10 NRVM preparations; asterisks denote significant differences among groups; ∗P <0.05 and ∗∗P<0.01; analysis using Ordinary one-way ANOVA and post hoc Holm Sidak’s multiple comparisons. D: Respiratory parameters such as basal respiration, ATP production, coupling efficiency, and maximal respiration were calculated from OCR data with and without PDE5i according to the mitochondrial stress test protocol. Bar equals mean ± SEM; each dot represents an individual patient serum-treated NRVM: n=12 healthy controls, n=8 SVHF samples +/− PDE5i; asterisks denote significant differences among groups; ∗P <0.05 and ∗∗P<0.01; analysis using Kruskal-Wallis test and post hoc Dunn’s multiple comparisons for basal respiration and ATP production, Brown-Forsythe and Welch ANOVA tests with post hoc Dunnett’s multiple comparisons for coupling efficiency, proton leak, and bioenergetic health index.

### 3.7 PDE5i Attenuates SV Serum-Induced Metabolite Changes in Cardiomyocytes

Given the observed alterations in metabolite profiles in response to SVHF serum-treatment, we assessed whether PDE5i therapy was sufficient to abrogate these changes (**Figure 6**). We determined that 12 of the 19 significantly differentially expressed metabolites (63%) were rescued with PDE5i treatment. The top 8 most abrogated metabolites in response to PDE5i were Glucose-6-Phosphate, Pyruvate, Fumarate, L-Carnitine, Decanoic Acid, Octadecatrienoic Acid, Bilirubin, and UDP-N-Acetyl-D-Glucosamine.

**Figure 6.**
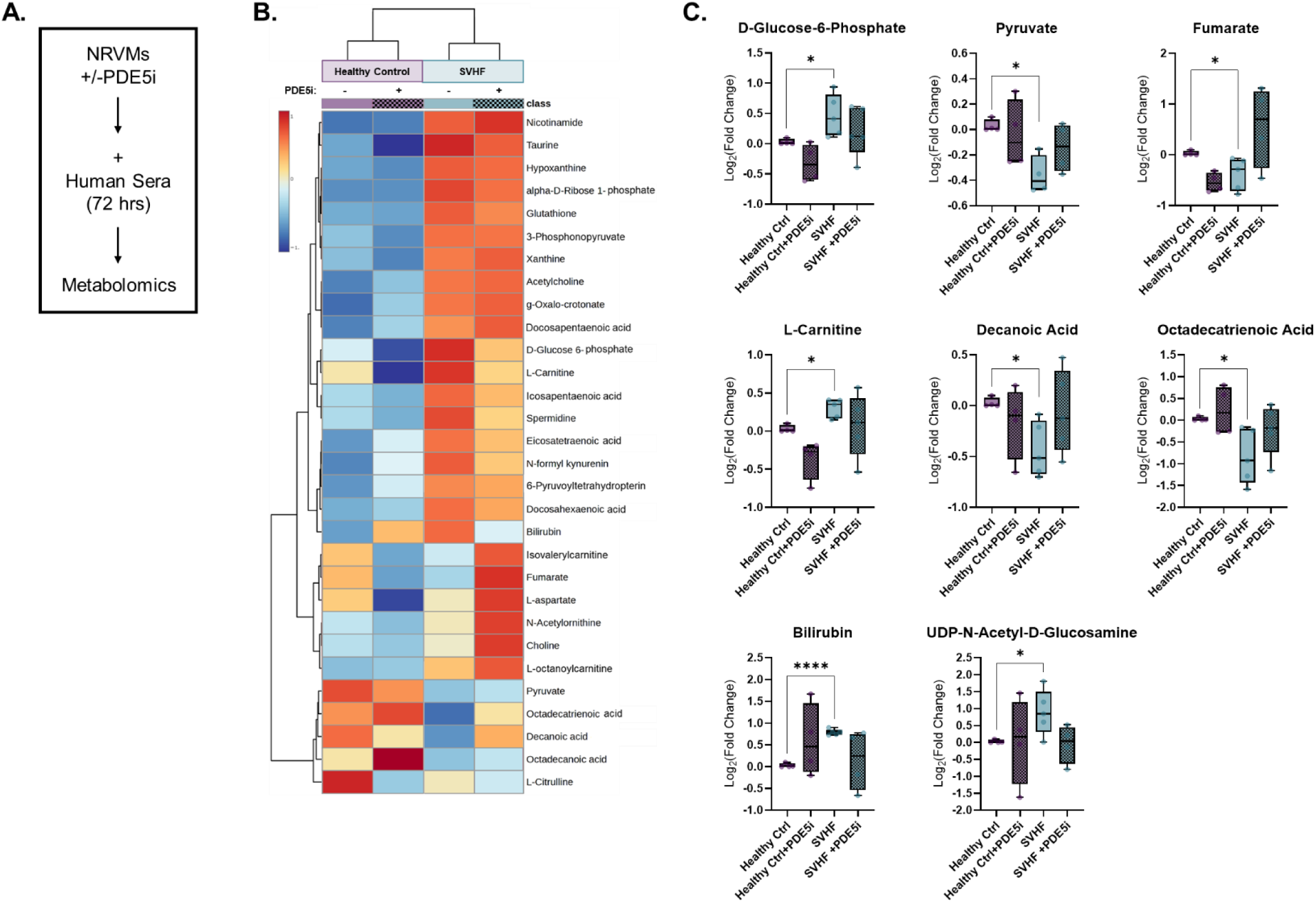
PDE5i Attenuates SV Serum-Induced Metabolite Changes in Cardiomyocytes *in vitro*. A: Mass spectrometry–based metabolomics analysis in serum-treated primary cardiomyocytes +/− PDE5i. B: Heatmap representation of the top 30 differentially expressed metabolites in serum-treated NRVMs +/− PDE5i. C: Log_2_ (Fold Change) of the top 8 metabolites attenuated with PDE5i, including D-Glucose-6-Phosphate, Pyruvate, Fumarate, L-Carnitine, Decanoic Acid, Octadecatrienoic Acid, Bilirubin, and UDP-N-Acetyl-D-Glucosamine; asterisks denote significant differences among groups; ∗P <0.05 and ∗∗∗∗P <0.0001; analysis using Brown-Forsythe and Welch ANOVA with post hoc Dunnett’s multiple comparisons. For all groups, average of n=4 (1 patient replicate) Healthy Controls and n=5 (2 patient replicates) failing SV –serum treated NRVMs across a total of 5 NRVM preparations.

### 3.8 PDE5i Does not Alter Relative Mitochondrial Copy Number in Cardiomyocytes

In order to investigate whether the metabolic changes seen in SVHF serum-treated NRVMs +/− PDE5i were due to alterations in the quantity of mitochondria, relative mitochondrial copy number was assessed via RT-qPCR (**Figure 7**). The expression of mitochondrial specific genes ND5 and CytB relative to nuclear B2M did not significantly differ between SVHF and Healthy Control serum-treated NRVMs +/− PDE5i.

**Figure 7.**
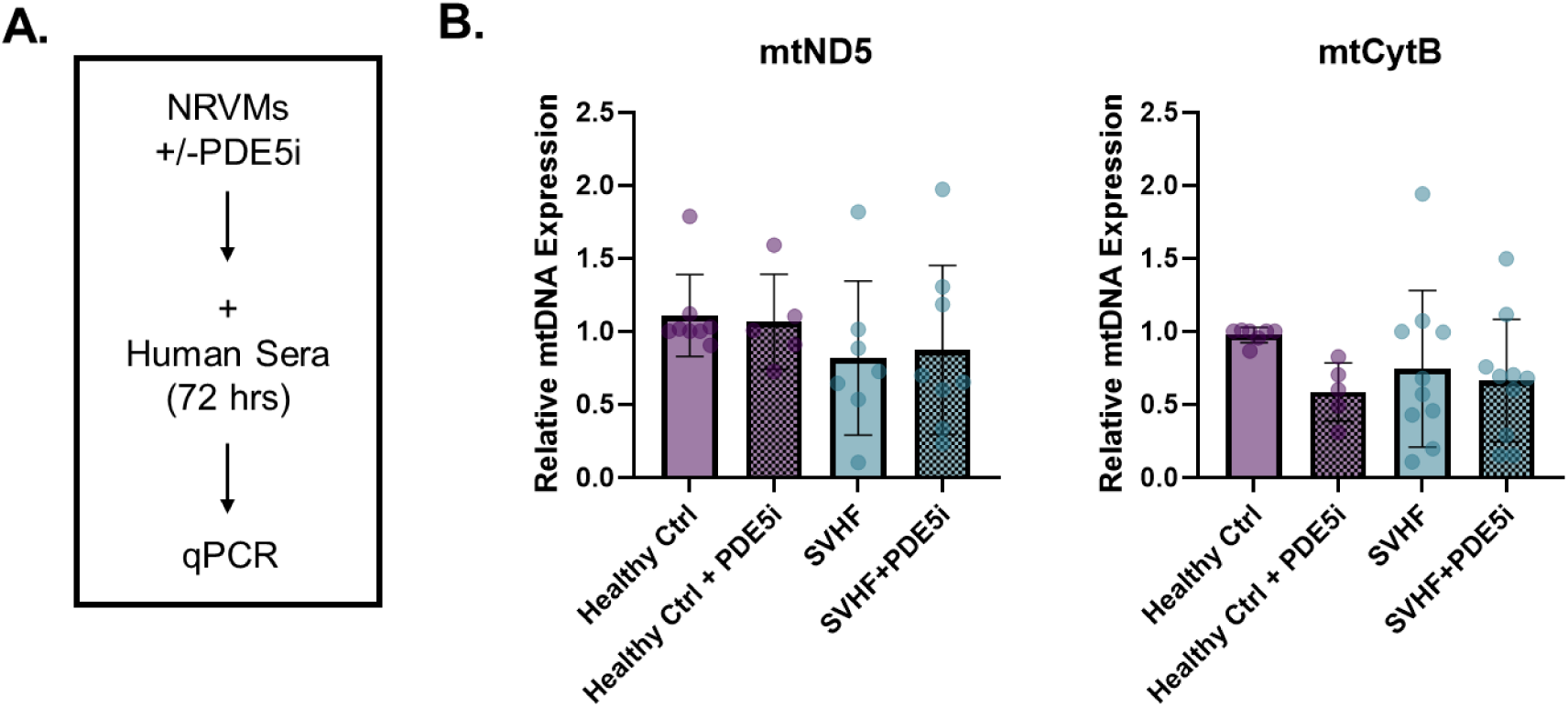
PDE5i Does not Alter Relative Mitochondrial Copy Number in Cardiomyocytes *in vitro*. A: DNA analysis of mitochondrially-encoded genes relative to nuclear-encoded genes from serum-treated NRVMs +/− PDE5i. B: Relative Fold Change of mitochondrial-encoded ND5 and CytB relative to nuclear B2M. Bar equals mean ± SEM; each dot represents an individual patient serum-treated NRVM: average of n=4 (3 patient replicates) Healthy Controls and n=5 (5 patient replicates) failing SV –serum treated NRVMs across a total of 6 NRVM preparations.

### 3.9 PDE5i Attenuates Impaired Fatty Acid Oxidation and CPT Activity in Cardiomyocytes

In order to evaluate the specific mechanisms by which SVHF sera and PDE5i therapy impact mitochondrial function, CPT I and CPT II enzymatic activity and palmitate oxidation were evaluated (**Figure 8**). Long-chain fatty acids (FAs) are well recognized as the preferred substrate for oxidative phosphorylation by cardiac mitochondria, and the mitochondrial CPT system is required for the delivery of long-chain FAs from the cytoplasm into the mitochondria for their subsequent β-oxidation. Compared to the Healthy Control serum-treated cardiomyocytes, SVHF treated cells had significantly impaired CPT I enzyme activity (P=0.0250), while CPT II enzyme activity had a non-statistically significant downward trend (P=0.12). Treatment with PDE5i therapy however, was sufficient to significantly improve CPT I and CPT II enzyme activity in SVHF treated cardiomyocytes (P=0.0459 and P=0.0320, respectively). Additionally, mitochondrial FAO in serum-treated primary cardiomyocytes +/− PDE5i was quantified using the addition of the long-chain fatty acid palmitate +/− Etomoxir (Eto), a CPT I inhibitor, using the same Cell Mito Stress Test in the Agilent Seahorse Bioanalyzer (**Figure 8**). Treatment of primary cardiomyocytes with failing SV patient sera resulted in significantly decreased basal respiration (P=0.0469), ATP Production (P=0.0422), and the overall Bioenergetic Health Index (P=0.0494). There were no differences in maximal respiration, spare respiratory capacity, proton leak, or non-mitochondrial oxygen consumption. PDE5i treatment significantly abrogated the SVHF serum induced changes in basal respiration and ATP production, and was sufficient to significantly decrease proton leak (P=0.0384). Moreover, PDE5i therapy significantly increased the overall bioenergetic health index (P=0.0059) in cardiomyocytes, which was abrogated with the addition of the CPT I inhibitor (P=0.0034).

**Figure 8.**
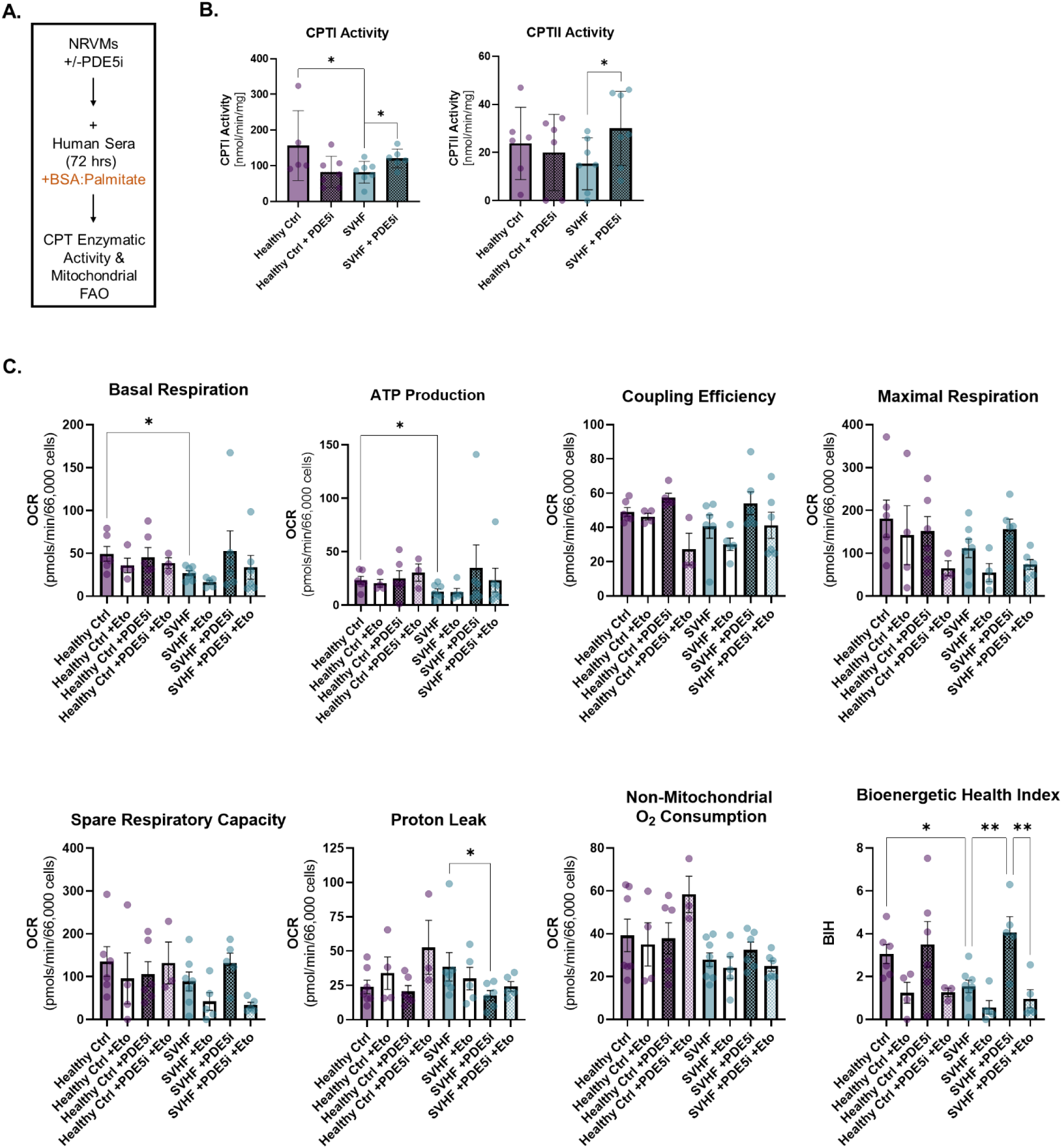
PDE5i Attenuates Impaired Fatty Acid Oxidation and CPT Activity in Cardiomyocytes *in vitro*. A: CPT enzymatic activity and mitochondrial FAO in serum-treated primary cardiomyocytes + the long-chain fatty acid palmitate, +/− PDE5i. B: Quantification (nmol/min/mg protein) of CPTI and CPTII enzymatic activity. Asterisk denotes significant difference among groups; ∗P <0.05; analysis using a Paired T-test; Bar equals mean ± SEM; each dot represents an individual patient serum-treated NRVM: n=6 healthy controls, n=7 SVHF serum-treated NRVM samples. C: Respiratory parameters such as basal respiration, ATP production, coupling efficiency, and maximal respiration were calculated from OCR data with and without PDE5i in the presence of BSA-conjugated palmitate according to mitochondrial stress test protocol. Bar equals mean ± SEM; each dot represents an individual patient serum-treated NRVM: n=6 healthy controls, n=4 healthy controls +eto, n=6 healthy controls +PDE5i, n=3 healthy controls +eto and PDE5i, n=8 SVHF samples, n=4 SVHF samples +eto, n=5 SVHF samples +PDE5i, n=4 SVHF samples +eto and PDE5i; asterisks denote significant differences among groups; ∗P <0.05 and ∗∗P<0.01; analysis using Brown-Forsythe and Welch ANOVA tests with post-hoc Dunnet’s multiple comparisons for basal respiration and ATP production, Kruskal-Wallis test and post-hoc Dunn’s multiple comparisons for proton leak, and ordinary one way ANOVA with post hoc Holm-Sidak’s multiple comparisons test for bioenergetic health index.

## 4. Discussion

Although increasing evidence suggests potential beneficial effects of PDE5i therapy on exercise capacity and ventricular function in the SV population, the specific mechanisms by which PDE5i elicits benefit remain unclear. We previously showed that *in vitro* treatment of primary cardiomyocytes with SVHF patient sera induces gene expression changes indicative of pathological myocardial remodeling, cardiac hypertrophy and cardiac dysfunction, which are abrogated by PDE5i treatment [21]. Utilizing the same model system, in the present study we investigated whether serum circulating factors promote metabolic dysfunction in SV, and if PDE5i therapy is sufficient to abrogate SVHF-serum induced pathological changes in cardiomyocyte mitochondrial bioenergetics.

Our prior data in SV myocardial tissue demonstrated that levels of cardiac cardiolipin, an inner mitochondrial membrane phospholipid that is critical for proper mitochondrial function, are significantly depleted in failing SV myocardium, with no significant change in relative mitochondrial copy number [27, 28]. Similarly, here we show that treatment of primary cardiomyocytes with failing SV patient sera significantly decreases total phospholipid levels, including a substantial decrease in the mitochondrial-specific phospholipid, CL. Therefore, SV serum circulating factors may play a role in modulating phospholipid dynamics in the SV heart, which may contribute to compromised membrane composition and function, including within mitochondria. PDE5i therapy however was not sufficient to attenuate CL remodeling, but did significantly increase PG levels in SVHF serum-treated cardiomyocytes. Mitochondria are capable of synthesizing several lipids autonomously, including PG, CL, and PE, and while mitochondrial PG is consistently low because it serves as an important precursor to CL biosynthesis, it can still play an important role in mitochondrial function. In fact, studies suggest that both PG and CL play an important role in mediating mitochondrial – nuclear crosstalk, and several PG species have been shown to interact with mitochondrial complex IV in the inner mitochondrial membrane[29]. Therefore, better understanding the impact of PDE5i on cardiomyocyte PG and further elucidating the function of PG in the context of SV are important. Notably, PG is the second most abundant phospholipid in the lung, playing a major role in surfactant function, therefore investigating the impact of PDE5i on lung phospholipid content in SV is also relevant[29].

In addition to alterations in the phospholipid milieu, we found that failing SV serum is also sufficient to induce significant changes in cardiomyocyte metabolomic profiles. Prior data demonstrated the failing SV heart is characterized by an altered abundance of TCA cycle metabolites, amino acid, acyl-CoAs, and acylcarnitines, suggesting global dysregulation of cardiac metabolism. Here, we demonstrate that SV serum circulating factors are sufficient to significantly increase levels of glucose-6-phosphate, and significantly decreased levels of pyruvate and fumarate, all suggestive of a metabolic switch from fatty acid β-oxidation. Importantly, PDE5i abrogated these potentially pathologic metabolite changes. Moreover, we found that SV serum significantly increased cardiomyocyte L-carnitine levels, which were abrogated with PDE5i therapy. L-carnitine plays an important role in cardiac energy metabolism, including interacting with the mitochondrial CPT system to facilitate the transport of long-chain fatty acids into the mitochondrial matrix for their subsequent oxidation. However, the implications of L-carnitine in HF remain paradoxical. While carnitine is used to treat some patients with HF there is little evidence to support the practice [30, 31]. The plasma levels of L-carnitine tend to be significantly elevated in patients with various cardiomyopathies, however this is inconsistent across studies. Interestingly, the cardiotoxic chemotherapy drug, Adriamycin, induces significant elevation of total plasma L-carnitine levels and concomitant inhibition of CPT enzymatic activity that was not rescued with exogenous L-carnitine supplementation[26]. Therefore, maintaining a homeostatic balance of L-carnitine levels in circulation and in cardiomyocytes is likely important for proper mitochondrial metabolic function in the heart.

The medium chain FA, decanoic acid, on the other hand, can be easily converted into ketones to serve as an alternative energy source, and several studies have shown its beneficial effects. For example, decanoic acid can improve mitochondrial function, decrease oxidative stress, and regulate blood lipid and glucose levels [32–34]. Here we show that failing SV sera significantly reduced levels of decanoic acid, which were recovered with PDE5i therapy, further suggesting a potential role for PDE5 in modulating metabolic substrate utilization in the heart. There is also increasing recognition that the O-linked attachment of N-acetyl-glucosamine (O-GlcNAc) on serine and threonine residues of proteins plays a role in modulating cell function and survival in the cardiovascular system. The levels of O-GlcNAc are regulated in part by glucose metabolism, and chronically elevated levels of O-GlcNAc may represent a common mechanism underlying the adverse effects of diabetes on cardiomyocyte, vascular, and endothelial cell function [35, 36]. Interestingly, failing SV sera were sufficient to significantly increase levels of O-GlcNAc in primary cardiomyocytes, and treatment with the PDE5i sildenafil attenuated this increase, suggesting yet another potential avenue by which PDE5i therapy may modulate cardiac metabolic function.

Given the critical need for a constant supply of energy in the beating heart, it is not surprising that many cardiovascular diseases involve disturbances in cardiac metabolism. SV serum metabolomic profiling for example, has identified significantly dysregulated circulating metabolites, including amino acids, energetic intermediates, and nucleotides indicative of metabolic remodeling that could be used for SV risk stratification, monitoring response to therapy, and even as novel targets of therapeutic intervention [37–39]. Additionally, our prior data identified that the SV RV myocardium is uniquely vulnerable to metabolic alterations, and the failing SV heart is typified by decreased mitochondrial oxidative phosphorylation and impaired cardiac energy generation. Similarly, here we show that treatment of primary cardiomyocytes with failing SV patient sera significantly impairs mitochondrial OXPHOS, illustrated by significantly increased ROS generation, and significantly decreased basal respiration, ATP production, and coupling efficiency – all contributing to an overall substantially decreased cardiomyocyte bioenergetic health index. The bioenergetic health index is a relatively new concept in metabolic research, and represents the composite mitochondrial profile for a selected cell type. A low bioenergetic health index is associated with a lower reserve capacity, low ATP-linked respiration and increased proton leak [40, 41]. Interestingly, we show that PDE5i therapy is sufficient to attenuate the SVHF serum-induced changes in ROS generation, ATP production, coupling efficiency, and substantially improve the overall bioenergetic health of cardiomyocytes.

In the healthy heart, long-chain FAs are well recognized as the preferred substrates for OXPHOS by cardiac mitochondria. The mitochondrial CPT system is responsible for the delivery of long-chain FAs from the cytoplasm into the mitochondria for their subsequent oxidation, and CPTI catalyzes the rate-limiting step. Our prior study demonstrated that the SV heart displays diminished mitochondrial CPTI and CPTII enzyme activity [28]. Functional analysis of the cardiomyocyte mitochondrial CPT system in response to SVHF serum-treatment demonstrated significantly decreased activity of CPTI. Intriguingly, PDE5i therapy significantly improved CPT activity, suggesting a novel role for PDE5 in modulating cardiomyocyte mitochondrial FA uptake and subsequent metabolic function. Correspondingly, we show that treatment of primary cardiomyocytes with failing SV patient sera significantly impairs mitochondrial FAO, illustrated by significantly decreased basal respiration, ATP production, and an overall substantially decreased cardiomyocyte bioenergetic health index in the presence of the LCFA palmitate. Importantly, we show that PDE5i therapy is sufficient to attenuate the SVHF serum-induced changes in FAO, including improved basal respiration, ATP production, proton leak, all contributing to substantial improvement of the overall bioenergetic health of cardiomyocytes. Importantly, inhibition of CPT I activity completely abrogated the positive effects of PDE5i therapy on cardiomyocyte bioenergetic health, suggesting PDE5i therapy modulates cardiomyocyte metabolism and substrate utilization via modulation of CPT activity.

### 4.1. Limitations

There are important limitations to this study. Studies of SV are challenged by small subject numbers due to the relative rarity of the disease. Because of this, we are not able to determine the influence of age, prior surgical procedures, or the temporal relationship of expression/activity changes in our findings. Secondary to the current widespread use of PDE inhibitors in the SV population, it is not possible for us to exclude all patients on a PDE5i, or to determine whether PDE5 upregulation is secondary to PDE3i (milrinone) treatment. It is important to note, however, that there was no significant difference in PDE5 activity or expression between SV patients based on treatment with the PDE5i sildenafil at the time of explant [42]. We recognize that comparisons of patients with SV at various stages of palliation may exhibit differences in systemic right ventricular volume load that may influence gene expression and metabolic function. Nevertheless, this study addresses important gaps in knowledge, by utilizing a novel model that poses no risk to children and provides valuable insights into the molecular pathways by which PDE5i therapy may modulate SV cardiac metabolism. This line of investigation deserves further study given the limited understanding of the mechanistic underpinnings of SV HF and the absence of evidence-based therapies to prevent or treat HF in this growing group of patients.

## 5. Conclusions

Together, these data suggest that SV serum circulating factors may potentiate progressive cardiac dysfunction and play a role in altering cardiomyocyte mitochondrial dysfunction and metabolic remodeling. Moreover, these data suggest PDE5i therapy contributes to beneficial cardiomyocyte metabolic remodeling in SV failure and as a result may have direct myocardial effects. Further investigation into exactly how PDE5i therapy alters mitochondrial substrate utilization and CPT activity in the SV population is warranted. Importantly, these data may have implications beyond the SV population, as a number of children with a wide range of heart and lung diseases are treated with PDE5i therapy.

## Acknowledgments

We would like to acknowledge the Blood Collection Team at University of Colorado Anschutz Medical Campus, including Dr. Stephanie Nakano, Andrew Shirley, Kaia Kinkel, Larissa Cussins, and Michaela Deck, our study coordinators Brett Li and Megyn Gordon for sample collection and study recruitment, and Meghan Williams for patient demographic curation.

---

**Supplemental Table S1.**
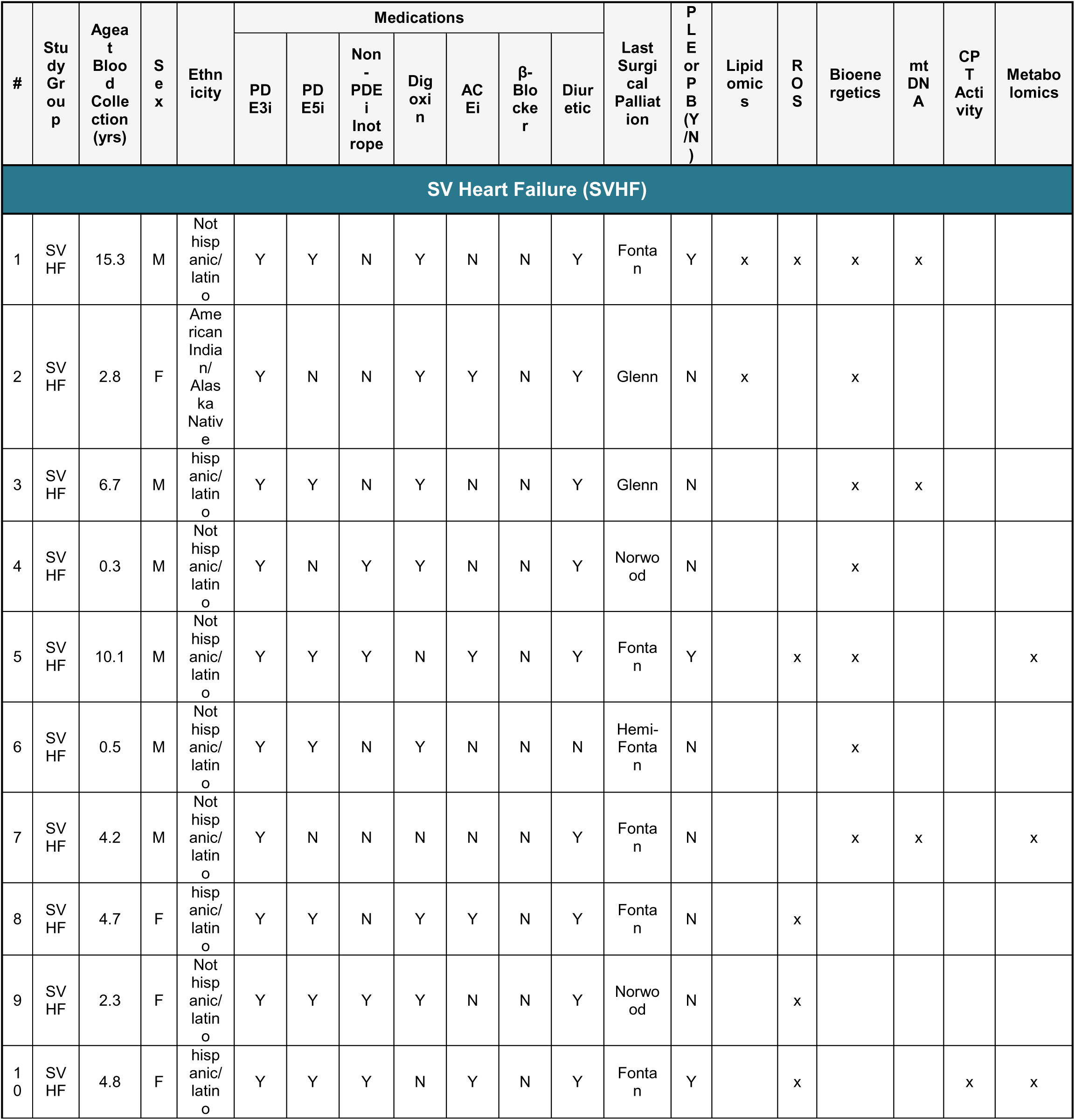
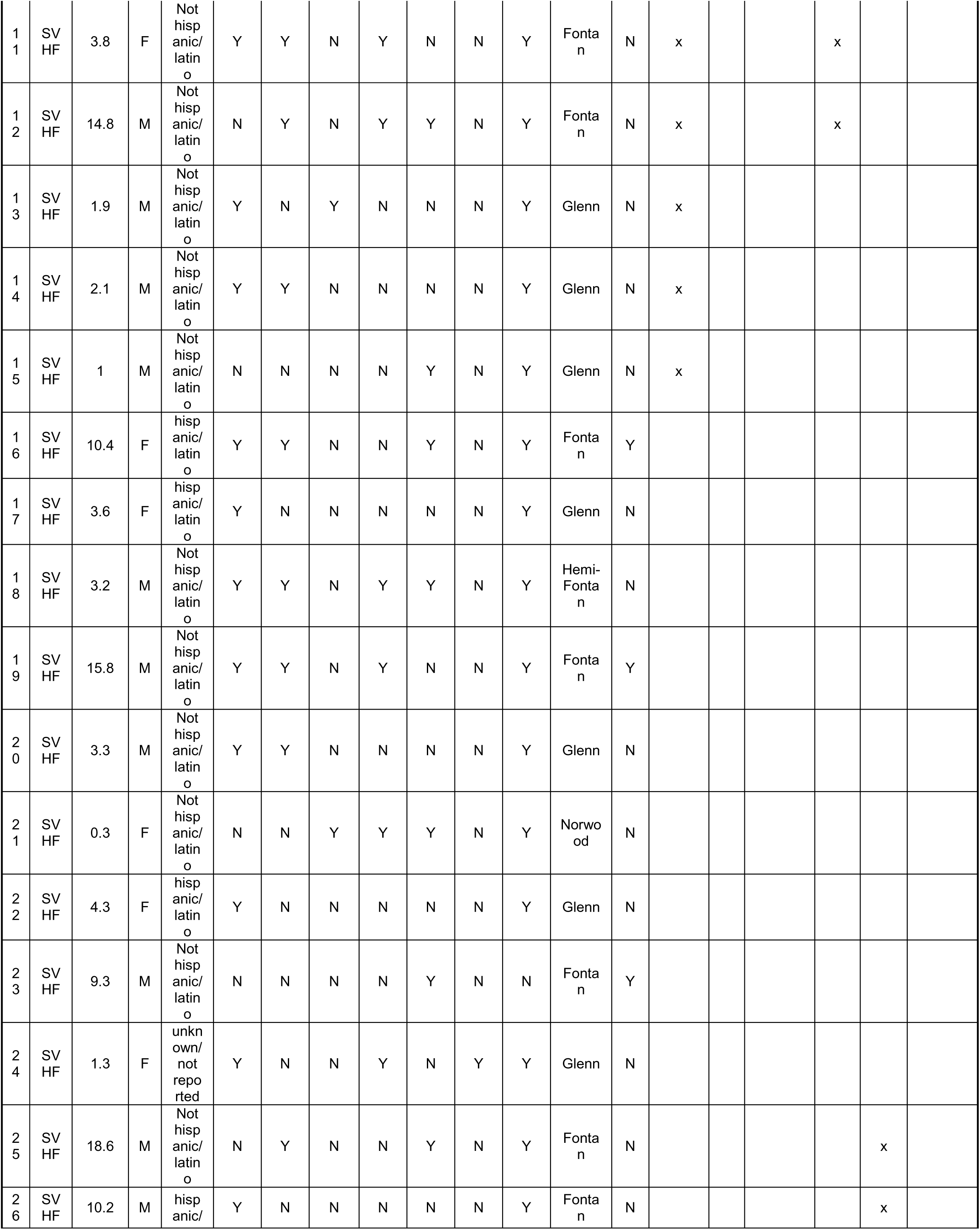
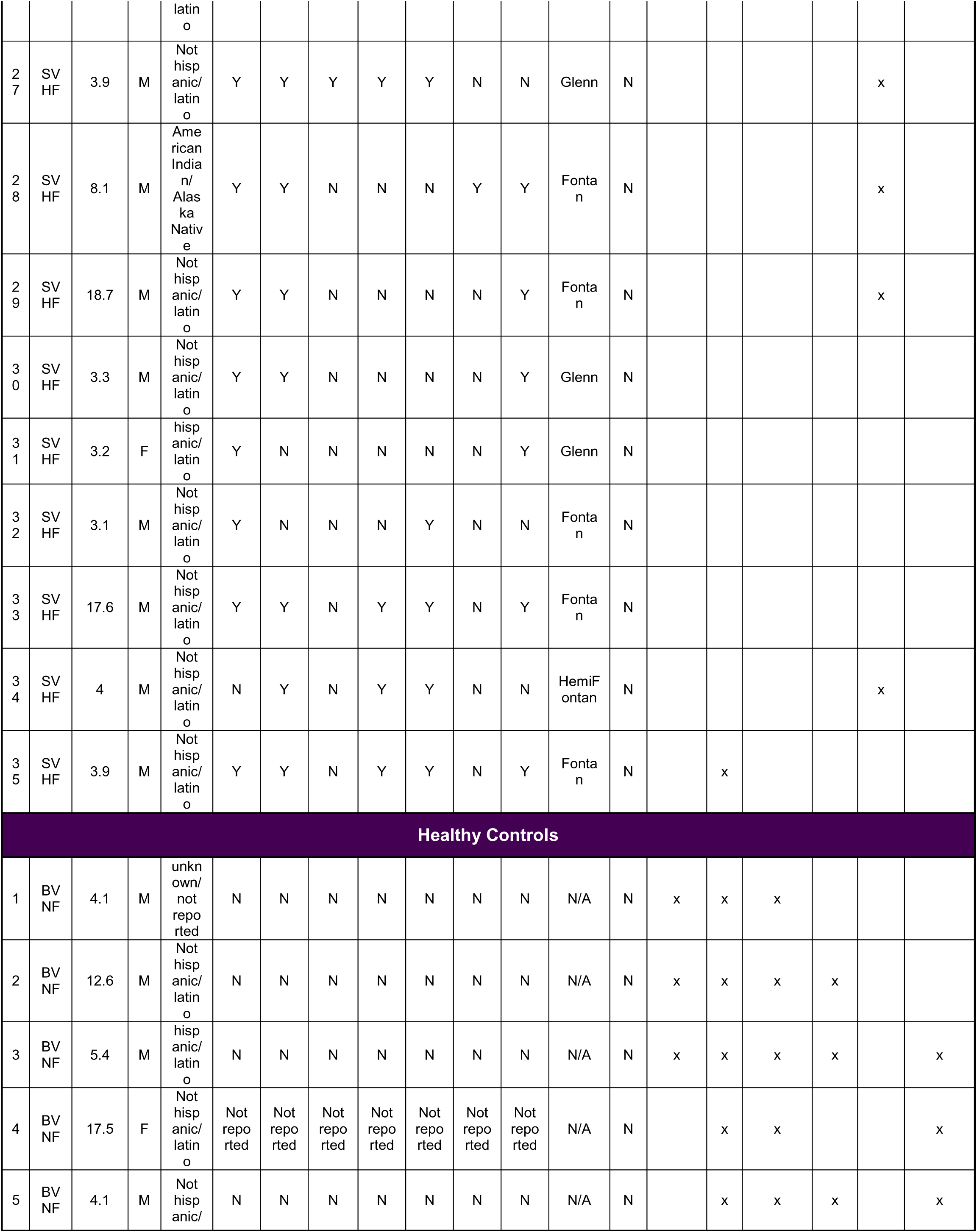
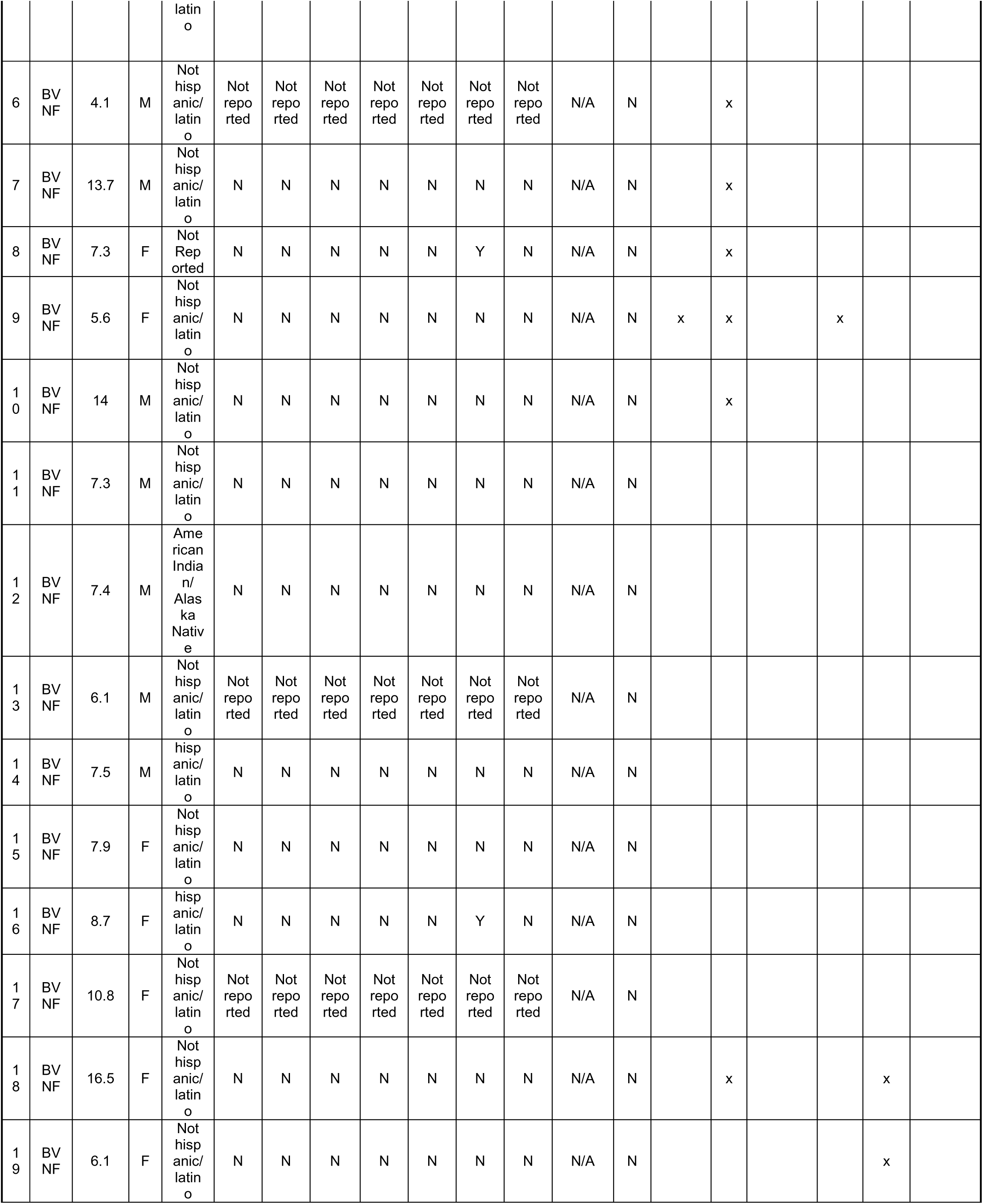
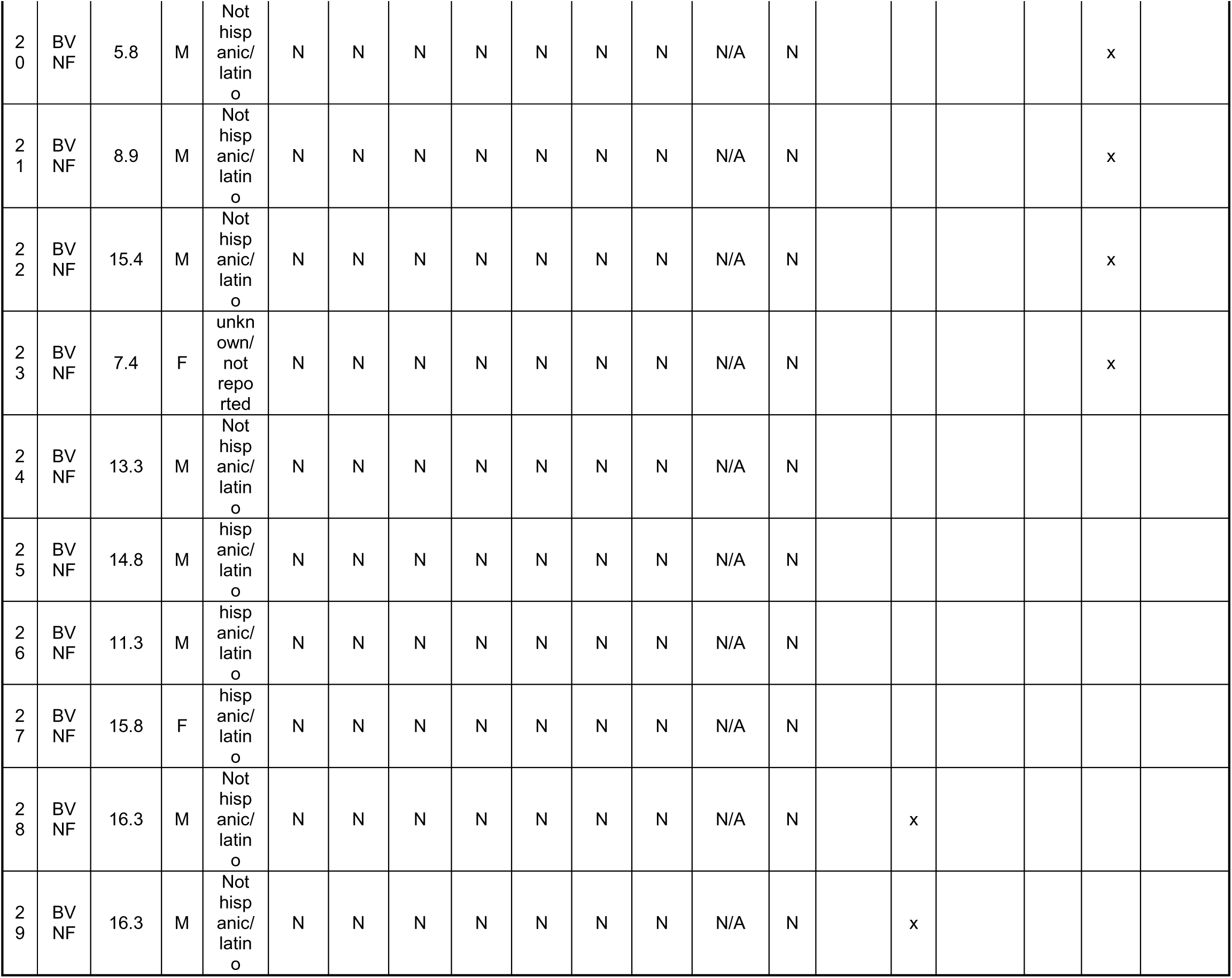
Patient Demographics. Complete demographics for all patients included in this study. ACEi = angiotensin-converting enzyme inhibitor, PDEi = phosphodiesterase inhibitor. *Non-PDEi Inotrope includes: dopamine, dobutamine, epinephrine, norepinephrine.

**Supplemental Table S2.** Major Cardiolipin Species by Molecular Weight. The m/z ratio and fatty acyl chain identification for each of the major CL species shown

| Major Cardiolipin Species by Molecular Weight |  |  |  |  |
| --- | --- | --- | --- | --- |
| Number of each Cardiolipin side chain constituent |  |  |  |  |
| Molecular Weight | *Palmitoleate 16:1* | *Linoleate 18:2* | *Oleate 18:1* | *Arachidonate 20:4* |
| 1422 | 1 | 3 |  |  |
| 1424 | 1 | 2 | 1 |  |
| 1448 |  | 4 |  |  |
| 1450 |  | 3 | 1 |  |
| 1472 |  | 3 |  | 1 |
| 1474 |  | 2 | 1 | 1 |
| 1496 |  | 2 |  | 2 |
| 1498 |  | 1 | 1 | 2 |

